# Plasticity and evolution of metabolic division of labour within families

**DOI:** 10.1101/2024.06.18.599519

**Authors:** EK Bladon, SM Hakala, RM Kilner, AC LeBoeuf

## Abstract

Fluids produced by parents for dependent young, such as milk or regurgitate, carry molecules that assist offspring with growth, immunity and digestion, allowing the metabolic burden of development to be shared between parents and offspring. We tested whether this division of metabolic labour changes plastically and evolves when offspring are experimentally deprived of their parents’ metabolic assistance. In the burying beetle *Nicrophorus vespilloides* parents deposit oral fluids on their carrion nest during pre-hatching care, and facultatively transfer fluids to larvae through oral trophallaxis as post-hatching care. We analysed the oral fluid proteomes of replicate experimental populations that had been evolving for 50 generations with or without post-hatching care, and which were then allowed to raise larvae with or without post-hatching care for one experimental generation. We found that parents and larvae plastically and evolutionarily adjusted the proteins in their oral fluids when we prevented post-hatching care. When reared in the absence of post-hatching care, larvae that evolved without post-hatching care were also more capable of consuming carrion proteins than larvae that had evolved with post-hatching care, and had higher survival. Our results suggest that metabolic division of labour within families is plastically modulated, and that the extent of socially modulated plasticity can evolve rapidly when social conditions change.

## Introduction

In species with parental care, the labour involved in protecting and provisioning offspring is divided between parents and their developing young^1^. Some duties are assumed by offspring from birth, such as digestion and other metabolic processes, and the path to offspring becoming fully independent of their parents is marked by the gradual acquisition of tasks that their parents initially undertook on their behalf, such as thermoregulation, immune defence, self-feeding and concealment from predators. The duration of this transition in division of labour varies between species and corresponds to the duration of postnatal parental care. Here we focus on the supply of parental care at the molecular level to investigate the extent to which this division of labour can vary plastically within species. We also use experimental evolution to investigate how rapidly offspring can evolve from being relatively dependent on their parents to having greater metabolic independence.

Our analyses focus on the molecular materials that are transferred from parents to offspring, and among offspring, because they provide a novel and precise way to quantify the metabolic division of labour. Socially transferred materials are seen across the animal kingdom and in diverse contexts (reviewed by ^2^). Within families, transfers occur not only between parents and offspring but also when siblings share materials (e.g.^3,4^). Socially transferred materials are typically a complex blend of molecules comprising the primary cargo e.g. nutritional macromolecules in the supply of care (e.g.^4^), and a secondary array of signalling molecules and metabolic shortcuts like proteins, RNAs and hormones, each of which can influence the short- and long-term physiology and behaviour of receivers^2^. These secondary components enable parents to assist offspring with vital functions like growth, immunity, digestion, and other metabolic processes, though at an energetic cost to parents (e.g.^5^). In other words, these molecules are a component of parental investment. Here we quantify the proteins in the fluids produced by both parents and offspring to assess the extent to which parents are sharing some of the metabolic burden of offspring development, both with each other and with their dependent young.

Our analyses focus on burying beetles *Nicrophorus vespilloides,* which provision offspring with both pre- and post-hatching care. In pre-hatching care, adults convert the dead body of a small vertebrate into an edible nest for their larvae. While they are making the nest, both adults transfer materials onto the flesh in oral and anal secretions, including molecules involved in immune defence and digestion^6,7^. The composition of these fluids varies plastically, according to the scale of microbial threat on the carrion^8,9^, the presence or absence of phoretic mites breeding on the carrion^10^ and in relation to contributions to lytic activity from other adults^6^.

During post-hatching care, adults transfer fluids by oral trophallaxis to larvae, whilst continuing to maintain the nest. Although adult oral fluids are known to enhance larval fitness^11,12,13^, it is not known whether they also transmit molecules with signalling or metabolic functions. The larvae secrete fluids orally and anally as well. These fluids are known to contain lysozymes^4^, but the larval fluids have not been analysed in any detail.

We compared the protein content of adult and offspring secretions collected from replicate experimental laboratory populations that had been allowed to evolve either with or without post-hatching care (*n* = 4 populations, details in ^14–19^). In two populations (Full Care), parents stayed with their offspring throughout their development to provide pre- and post-hatching care. In the two other populations (No Care), we removed parents when the nest was completed but before the larvae hatched, so offspring in these populations received no post-hatching care at all. These two regimes of care were imposed in the same way on each population for over 50 generations in total. We have previously reported that the No Care populations rapidly evolved and adapted to the absence of post-hatching care^14–21^. After 13 generations, parents in the No Care populations had evolved to ‘frontload’ parental care and put more effort into nest-preparation than Full Care parents^15^. In addition, larvae in the No Care populations had evolved to help each other gain mass in the absence of parental care, through mechanisms that have yet to be identified^18^.

To test whether the division of labour between parents and offspring had evolved divergently in the No Care and Full Care populations, we used nano-liquid chromatography tandem mass spectrometry (proteomics) to analyse the protein composition of burying beetle adult and larval oral fluids after 48 generations of experimental evolution. We allowed members of the No Care and Full Care populations to each experience Full Care conditions for two generations (49 and 50) prior to sampling at generation 51 to remove any parental or grandparental early-life effects. This design allowed us to investigate the degree of larval dependence on post-hatching care, the extent to which adults and larvae modulate the protein content of their fluids in relation to each other, and whether the extent of this plasticity can evolve (Figure 1).

**Figure 1:**
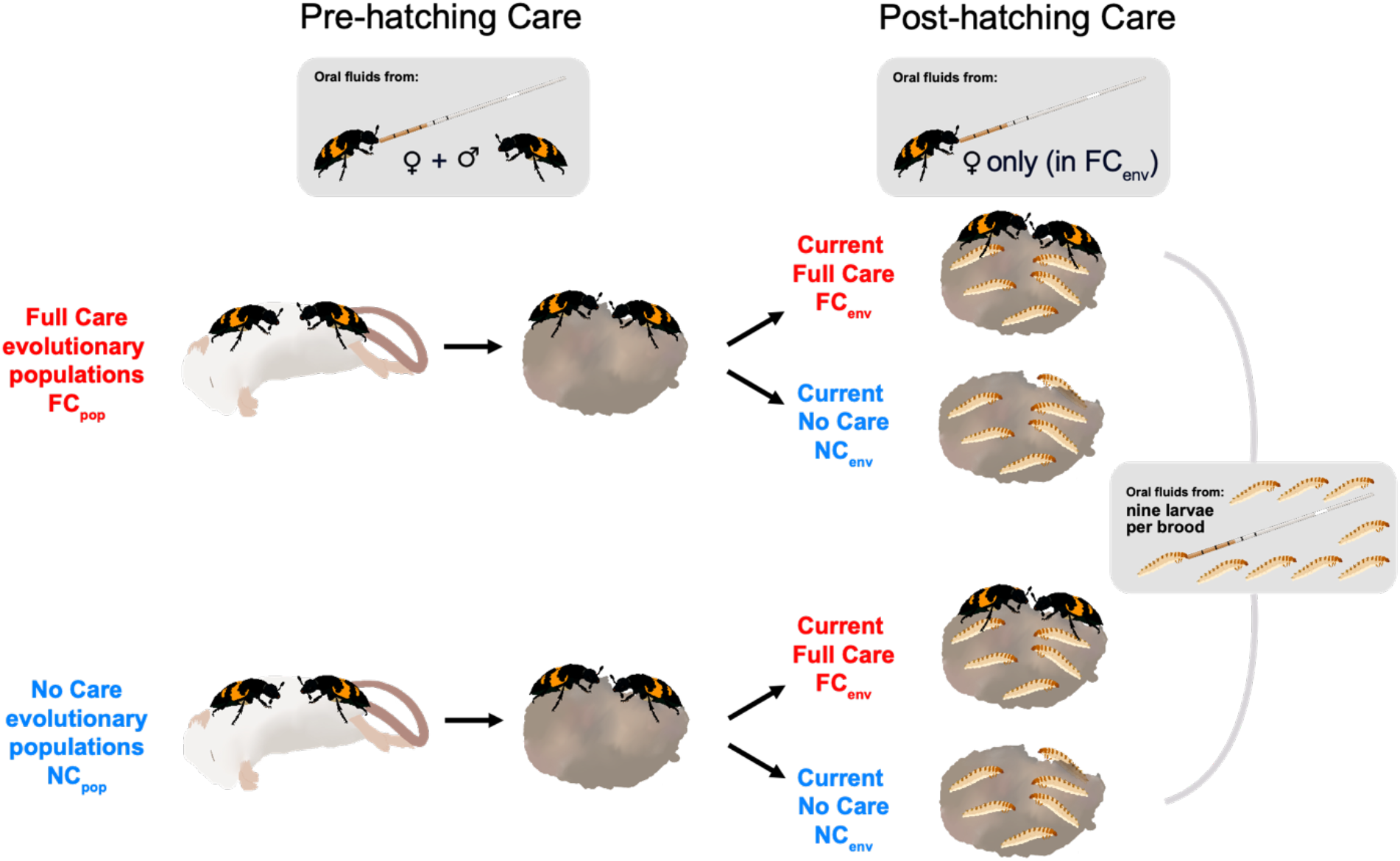
Experimental design of the oral fluid sampling. The pre-hatching care sampling time was 53 h after parents were paired with a mouse carcass and the post-hatching time sampling time was at the 3^rd^ larval instar. See supplementary video 1 for fluid collection technique.

## Results

### Larvae that evolved with parental care consumed fewer mouse proteins without parents

Oral fluids in all our samples contained a mixture of beetle-derived and mouse-derived proteins, generally with a higher proportion of beetle-relative to mouse-produced proteins, when considering the total protein intensity assigned to each species (Figure 2A, *t* = 6.6681, *P* <0.0001). For larvae, we could not distinguish whether the beetle proteins found in larval regurgitate came from parents, via the carcass, or were produced by the larvae themselves. Larvae generally had higher amounts of mouse protein in their oral fluid than did adults (Figure 2A, supplementary table 1), likely because they were feeding constantly on the carcass. However, larvae that *evolved with* full post-hatching care but were *reared without* post-hatching care had the lowest mouse-protein proportion (Figure 2A, supplementary table 1), and also the lowest absolute mouse-protein intensities (on average 6×10^11^, compared to the three other larvae types that range between 14-17×10^11^) though this effect seems to be driven by only one of the two experimental evolution replicate blocks (Figure 2A). This suggests that these larvae had a greater dependence on post-hatching care for access to exogenous resources, which might account for their lower survival when deprived of post-hatching care.

**Figure 2:**
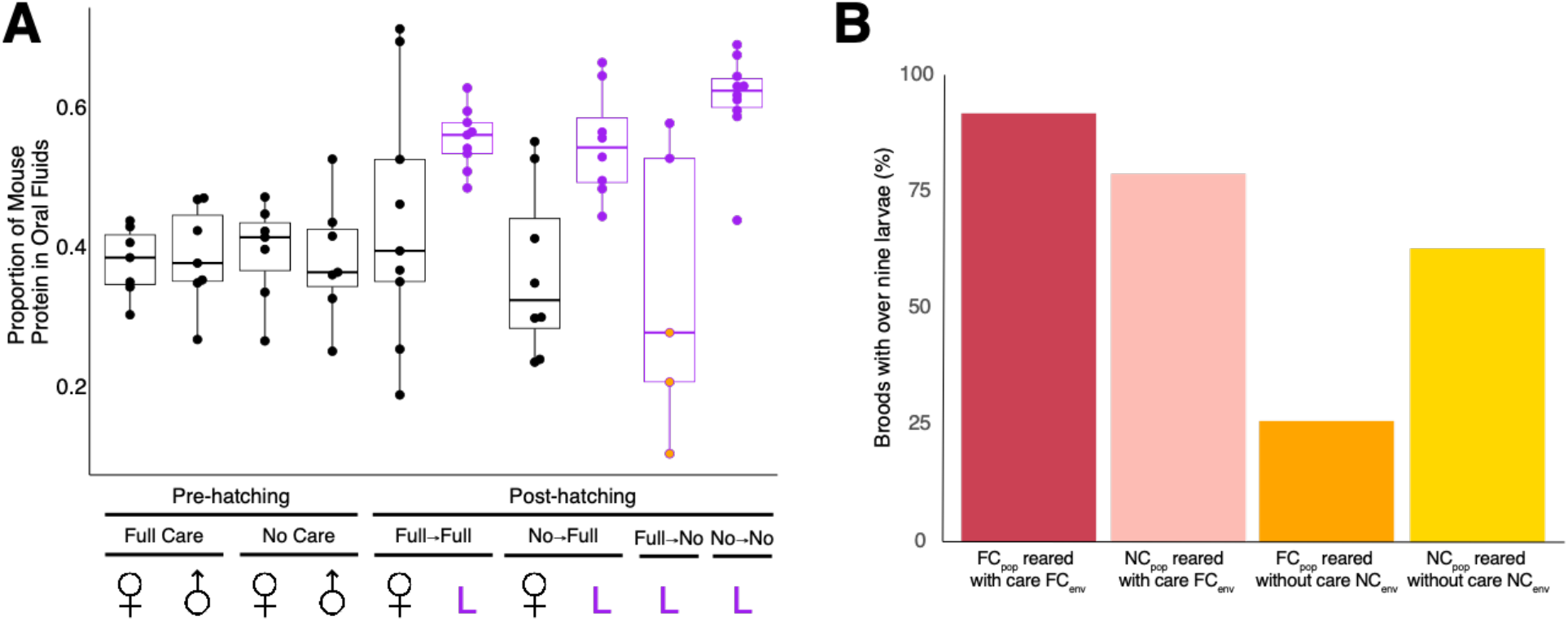
Impact of evolutionary population and current care regime on larval fitness. A) The proportion of mouse-derived proteins in beetle oral fluids collected from females, males and larvae (L). For each type of sample, we show the proportion of intensity assigned to mouse proteins. For larval samples the first word of the treatment name refers to the level of care they evolved with, and the second word refers to the level of care they experienced in the experimental generation (‘Full’ = Full Care, ‘No’ = No Care). Purple indicates larval regurgitate and black adult regurgitate. The orange-filled points in the Full Care to No Care larval regurgitate indicate that these three points were from one of the two experimental replicate populations. This was the only condition where the two replicates differed. Bold horizontal lines on the box plots represent the median values. Whiskers extend to the farthest data point which is no more than 1.5 times the interquartile range from the box. B) The percentage of broods that had nine or greater larvae surviving to third instar.

This variation in dependence is also apparent when assessing brood survival across the conditions. Only 26% of Full Care population broods reared with no post-hatching care had at least nine surviving larvae, compared to 92% of Full Care population broods reared with full post-hatching care, 63% of No Care population broods reared with no post-hatching care and 79% No Care population broods reared with full post-hatching care (Figure 2B).

### Beetle proteins in oral fluids are related to digestion, immunity and growth

To better understand the potential functions of the beetle-derived proteins in these oral fluids, we analysed the 200 beetle-derived proteins identified in the samples (supplementary table 2). We found *Drosophila melanogaster* orthologues for each beetle protein where possible, and classified the beetle proteins into 38 that were growth-related (e.g. larval serum proteins, development-related proteins), 82 digestion-related (e.g. involved in proteolysis), 28 immunity-related (e.g. lectins) and 14 communication-related (e.g. odorant-binding protein) (Figure 3). The most abundant protein, a serine-type endopeptidase, constituted 8-19% of all beetle-derived protein on average per sample type, while the rank abundances of other proteins diminished rapidly, consistent with other socially transferred materials^22,23^ (Figure 3). Fifty-six proteins could not be classified into these four main groups, including four proteins with orthologues in *Drosophila* ejaculate, several structural and housekeeping proteins (Hsp70, ribosomal proteins), several with unknown functions, and 13 lineage-specific, novel uncharacterized proteins. Oral fluid samples contained different proteins from the two control haemolymph samples, showing there was no notable haemolymph contamination in the oral fluids sampled (supplementary figure 1).

**Figure 3:**
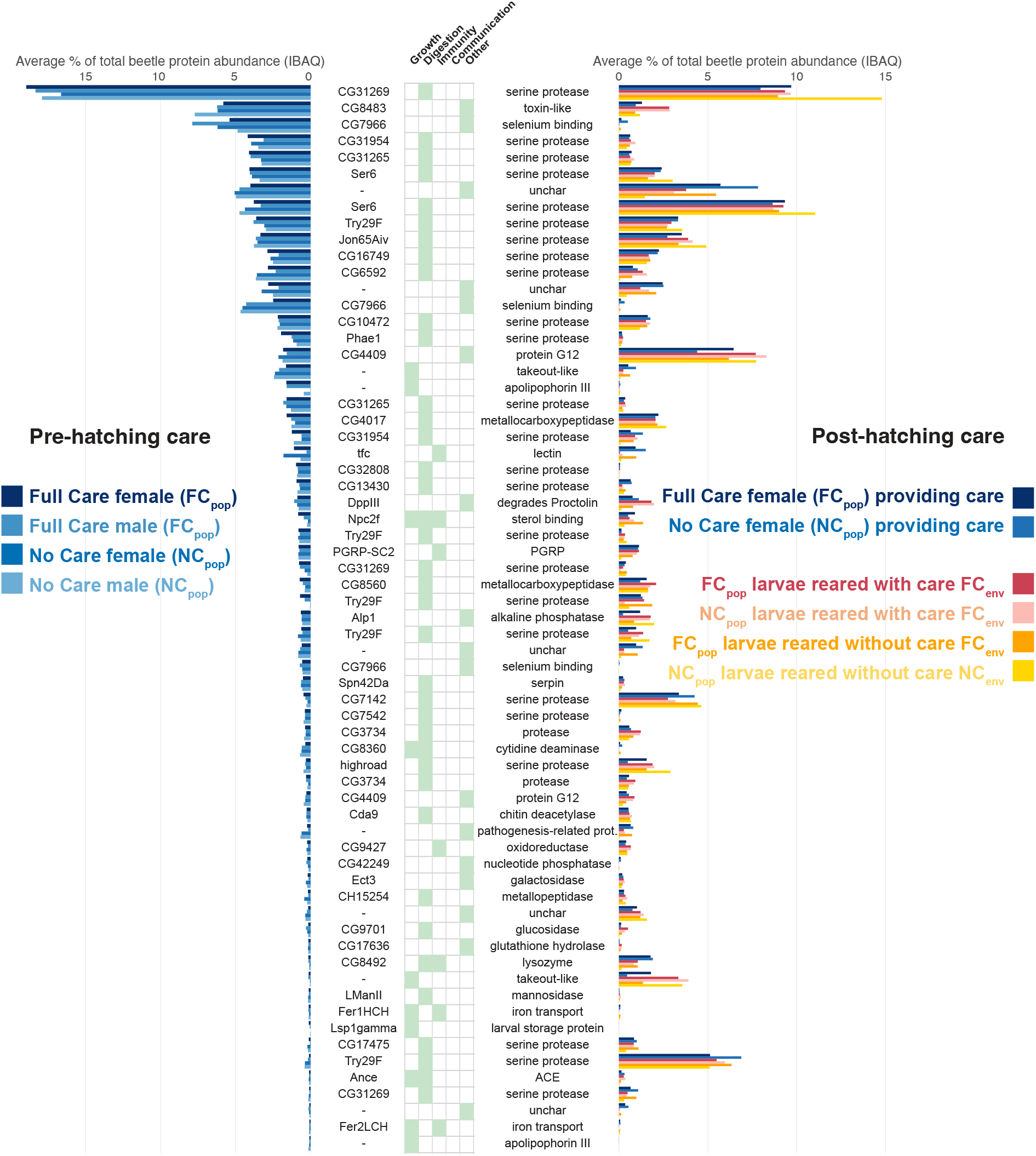
The 65 most abundant proteins in oral fluids produced by parents and offspring, in order of abundance in the Full Care females sampled pre-hatching. Bars on the left-hand side show samples taken from parents during carcass preparation (pre-hatching care). Bars on the right-hand side show samples taken from mothers during post hatching care (blues) and from 3rd instar larvae (post-hatching care). Proteins are annotated with their *Drosophila melanogaster* orthologue genes, our function classification, and a summary of available annotations. Abundance is shown as the average percentage of the total iBAQ assigned to that protein, calculated separately for each sample type. Proteins that do not have *D. melanogaster* orthologues are annotated with a hyphen. Protein accession numbers are presented in detail in supplementary table 2.

To further analyse the types of metabolic function the oral fluid proteins could be facilitating, we carried out a gene set enrichment analysis of gene ontology terms, pathways and protein-protein interaction networks of the oral fluid proteins’ *Drosophila melanogaster* orthologues. The orthologues of the 65 most abundant proteins in adult and larval burying beetle oral fluids, pre- or post-hatching, or the proteins significantly differing in any of our analyses, primarily have molecular functions involving peptidase and hydrolase activities, e.g. “peptidase activity” (FDR < 1e-18, supplementary table 3), indicating that these are primarily digestive fluids.

### Parents from the No Care and Full Care evolutionary populations differ in their oral fluid proteins during pre-hatching care

We have previously found that parents from the No Care populations evolved to frontload parental care by putting greater effort into nest preparation^15^. We therefore predicted that during nest preparation, these No Care parents would be more likely to have proteins with digestive, immune or larval growth functions in their oral fluids.

We found that during pre-hatching care the oral fluid proteomes of the two types of evolutionary population clustered separately by their first two principal components (Figure 4A), suggesting that they had indeed followed different evolutionary trajectories. However, the populations diverged in ways we did not predict. Six proteins differed in abundance between the populations during pre-hatching care (Figure 4C). Five proteins (three growth-related proteins: a larval serum protein, apolipophorin, and a sterol-binding protein, and two digestion-related endopeptidases) were more abundant in the Full Care populations – contrary to our prediction (Figure 4, supplementary table 4). The only protein that was more abundant in the No Care parents pre-hatching is potentially a thaumatin-like anti-fungal protein without a Drosophila orthologue^24^.

**Figure 4:**
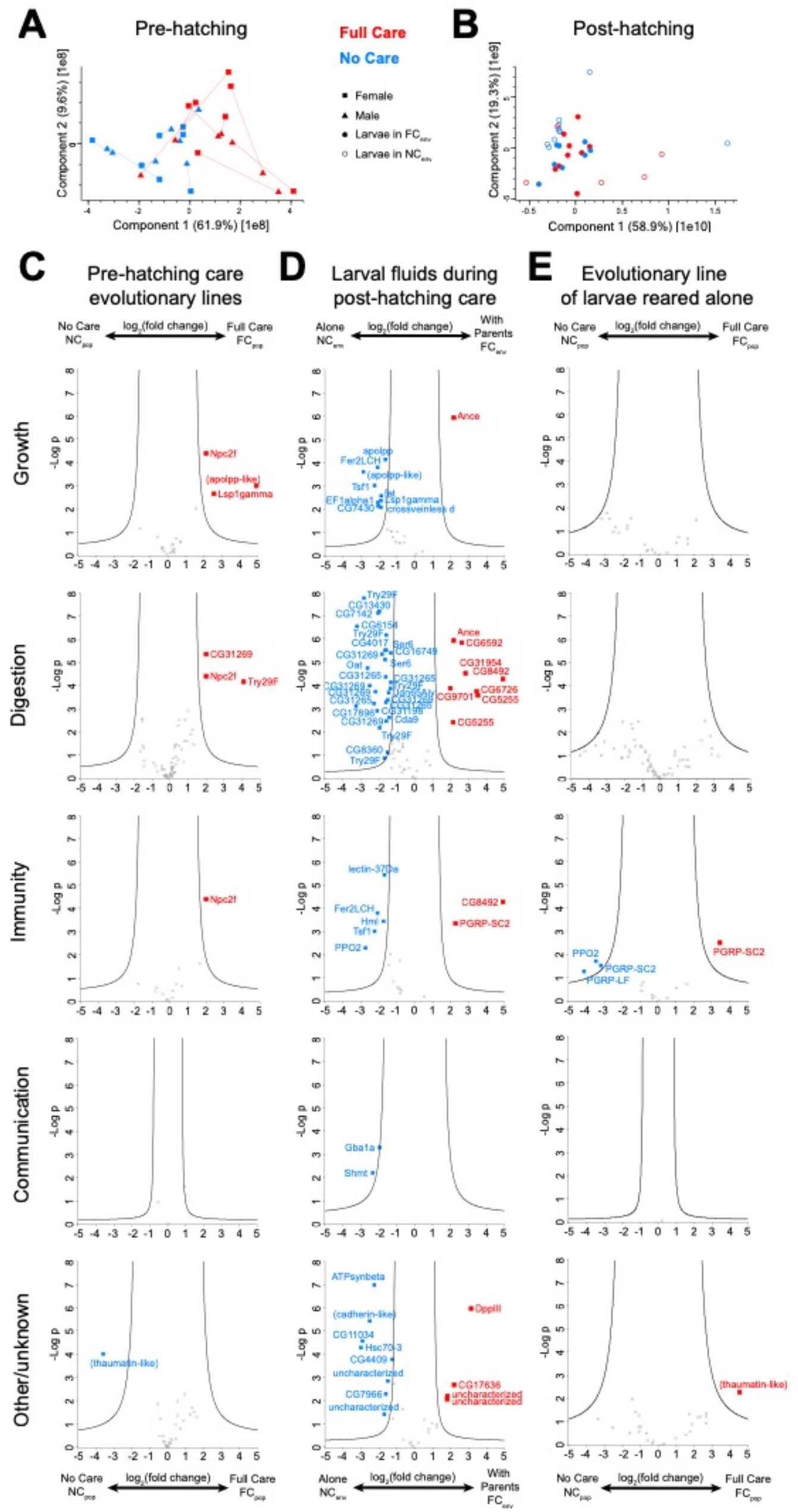
Protein composition shifts with evolutionary care environment and current care environment. Principal components analyses for A) male and female oral fluids at the pre-hatching time point with pairs connected with lines, B) larval oral fluids with and without post-hatching care in the experimental generation (all treatments). Red = Full Care evolutionary populations, blue = No Care evolutionary populations. Squares = females, triangles = males, full circle = offspring with post-hatching care, open circle = offspring with no post-hatching care. Volcano plots (C-E) showing the fold-changes and negative log-p values of the differing protein abundances by sample type (column) and protein type (row), annotated with their *Drosophila melanogaster* gene orthologues. Where no *D. melanogaster* orthologue was available, proteins are listed as uncharacterised, or with a BLAST-derived annotation in parenthesis. C) The oral fluid proteins found in pre-hatching care fluids that differ in abundance across evolutionary populations, where proteins more abundant in No Care are in blue (NC_pop_) and Full Care in red (FC_pop_). D) Larval fluids during post-hatching care, differing by current care regime (FC_env_ or NC_env_). E) Fluids of larvae reared without post-hatching care (NC_env_), with protein abundances differing between evolutionary populations (FC_pop_ or NC_pop_). Full results in supplementary table 4.

Next, we tested the extent to which males and females within pairs adjusted the protein content of their oral fluids in response to each other during pre-hatching care. Overall, proteins were similarly abundant in the two sexes (supplementary table 4). However, in the PCA (Figure 4A), males and females from the No Care populations clustered together more closely within pairs than their counterparts from the Full Care populations. The overall Euclidean distance separating members of the same pair was significantly smaller for No Care pairs than it was for Full Care pairs (*t* = −3.356, *P* = 0.001, Figure 4, Supplementary table 4), indicating a greater similarity in the protein profiles produced by two No Care partners. Within Full Care pairs, the Euclidean distance between males and females was not significantly different from the distance between ‘pairs’ that we randomly generated *post-hoc* from the Full Care dataset (*t* = −1.419, *P* = 0.16, supplementary table 5).

### Both current and evolutionary care regimes impact larval oral secretions

We found that the larval oral fluids differed in their protein composition between the four treatments (NC_POP_NC_ENV_, NC_POP_FC_ENV_, FC_POP_NC_ENV_, FC_POP_FC_ENV_, Figure 3 and Figure 4). However, most differences were due to whether or not larvae received care in the current generation, rather than due to their evolutionary population of origin (Figure 4). Sixty-two proteins differed significantly in abundance depending on whether or not larvae were currently receiving post-hatching care (Figure 4D, supplementary table 4) and 49 of them were more abundant when the parents were absent (24% of all proteins annotated as growth, 33% of digestion, 18% of immunity, 14% of communication and 14% of other), suggesting that larvae can compensate when parents do not supply these proteins. We detected some evolved divergences between No Care and Full Care populations, but only when larvae developed without post-hatching care (Figure 4E). Five proteins differed significantly in abundance between larval fluids across evolutionary populations in the No Care environment (NC_ENV_), with three immune-related proteins being higher in the No Care populations (prophenoloxidase and two proteoglycan recognition proteins, Figure 4E, supplementary table 4) and two being higher in the Full Care populations (a different proteoglycan recognition protein and a thaumatin-like protein, Figure 4E, Supplementary table 4). When larvae received post-hatching care (FC_ENV_), only cytidine deaminase (CG8360, supplementary table 4), a key protein for pyrimidine salvaging, was produced more abundantly by larvae in the No Care evolutionary populations.

### Oral protein secretions from mothers of the evolutionary populations do not differ during post-hatching care

The abundance of proteins in the oral fluids of mothers from the Full Care evolutionary populations did not differ significantly from those in No Care evolutionary populations (supplementary table 4). All maternal samples formed a single mixed cluster in the PCA analysis (supplementary figure 2B), and the coefficient of variation of the protein abundances did not significantly differ between Full Care and No Care females (*t* = −1.876, *P* = 0.061). We conclude that the presence of offspring is such a strong social cue for the mothers that they always provide post-hatching metabolic resources for their offspring, regardless of their recent evolutionary history of care, and even when that care is somewhat less effective (see survival data above).

### Do mothers and larvae fine-tune their oral protein secretions in response to each other?

We found no evidence that maternal and larval proteomes either matched or complemented each other. Pooling data from both types of evolutionary population, we found that the Euclidean distance between the proteomes of female-larvae pairs did not differ between true mother-offspring pairs and pairs randomly generated *post-hoc* (*t* = −0.674, *P* = 0.501, supplementary table 5). We also found that the Euclidean distance between mother-offspring pairs did not differ significantly between the No Care and Full Care populations (*t* = −1.528, *P* = 0.128, supplementary table 5).

## Discussion

To understand how metabolic labour is divided within families, we have described a molecular form of parental care in detail by characterising and quantifying the proteins in the oral fluids produced and shared by the different members of burying beetle *N. vespilloides* families. By combining evolving populations with manipulations of care provided in one generation, we have clarified how this metabolic division of labour changes plastically and over short evolutionary times.

### Oral fluids enable metabolic division of labour

An overwhelming majority of the proteins that we found in burying beetle oral fluids had *Drosophila* orthologues with metabolism-related functions in growth, immunity and digestion, suggesting that parents share some of the metabolic burden of development with their offspring by producing enzymes that will help them develop. If a parent produces a digestion enzyme, or a pathogen recognition protein in abundance that their offspring can use, then offspring do not need to expend the energy and resources to produce this protein themselves and can simply benefit from its action. The social transfer of proteins that function in metabolism therefore provides a mechanism for the division of metabolic labour between family members. For example, adult oral fluids produced before larvae hatched contained storage proteins such as the hexamerin Lsp1gamma and Apolipophorin III. Both of these have been implicated in within-individual hormone, lipid and amino-acid traffic^23,25–27^ and their orthologues are also present in oral care fluids of the ant *Camponotus floridanus*^23^. Sharing such consolidated resources, which result from many enzymatic steps of catabolism and anabolism, may provide offspring with significant metabolic shortcuts. Similarly, Niemann–Pick C proteins (NPC2 proteins) were also detected in beetle oral fluids. They have previously been found in ant regurgitated fluids and have been implicated in insects in sterol binding and ecdysteroid biosynthesis^28,29^. They could function directly to promote healthy larval development or act as carriers for other molecular cargo^30^.

Although many of the oral fluid proteins had metabolic functions, some could serve a signalling or carrier function too. All the oral fluids we sampled contained serpins in abundance, which function as serine-protease inhibitors and could act on the numerous serine proteases in the fluid. In vertebrates, serpins have also evolved into carrier proteins for important cargo^31^. Numerous proteins we found could act in cargo transport (serpins, NPC proteins and odorant binding proteins) and could also play a role in signalling the location of the carrion nest to newly hatched beetle larvae, who hatch in the soil surrounding the carrion and must then crawl to the nest.

In a clear link with previous literature on burying beetle parent care, we found two takeout-like proteins in the oral fluids, one high in pre-hatching caring adults (XP_017777889) and one produced by both adults and larvae during the post-hatching phase (XP_017777894). Takeout genes were found to be differentially expressed in the transition to parenting in *N. vespilloides* ^20,32,33^ and *N. orbicollis*^34^. While neither of the proteins observed here are the precise genes or orthologues previously implicated in *vespilloides* or *orbicollis* they are within the same gene family which has sustained several duplications in this lineage relative to *Drosophila*. These previous gene expression screens also implicated storage proteins like Apolipophorin-III in parental care^34^. Together with our findings, these results strongly suggest that the expression of parental care involves a shift to physiologically provisioning young with parent-derived molecules.

### Evolved change in the expression of oral fluid proteins before hatching in parents

We expected to find that parents of No Care populations secreted more proteins in their oral fluids during nest preparation than Full Care parents, since we have previously shown that No Care parents invest more effort in other aspects of nest preparation than Full Care parents^15^. However, we found the opposite pattern, with Full Care parents producing more of some specific proteins than No Care parents. One potential adaptive explanation centres on a putative trade-off within parents between the effort they expend on shaping the carrion nest and the effort they spend on producing oral proteins. Perhaps in the No Care populations, preparing a rounder nest more rapidly yields a greater fitness return for offspring than adding certain proteins to the surface of the flesh – whereas the reverse is true in the Full Care populations. It would be interesting in future work to test whether such a trade-off indeed exists, and whether it can be flexibly modulated. An alternative interpretation is that a sustained absence of parental care over the generations in the No Care populations yielded individuals of such inferior quality that they were incapable of producing more proteins. This interpretation seems less likely, though, because we eliminated parental and grandparental effects as much as possible by putting all experimental individuals through two Full Care common garden generations before sampling their oral fluids.

Although No Care parents did not evolve to produce more oral proteins before hatching than Full Care parents, care-giving partners from the No Care populations matched the protein content of their fluids to a greater extent than individuals from the Full Care populations. Task specialisation throughout parental care is seen in natural populations of burying beetles, where males specialise in pre-hatching care^35,36^, females stay with the larvae for longer^14^, transfer more fluids mouth-to-mouth^35,36^ and produce higher levels of antimicrobial content in their anal exudates^6^. The data we present here suggest that by eliminating post-hatching care in the No Care populations, we also started to erode the extent of task specialisation by each sex.

### Evolutionary reliance on post-hatching care

After hatching, we found no difference in the proteins secreted orally by No Care and Full Care mothers when they were exposed to a brood. This is consistent with other assessments we have made of the No Care females’ capacity to provide care which, unlike male care, remained unchanged despite not being expressed for many generations^14^. By contrast, larvae deprived of post-hatching care showed a greater change in their oral secretions which, for larvae from Full Care populations, was associated with a decrease in survival. This suggests that Full Care larvae are more dependent than are No Care larvae on metabolic assistance from their parents during development ^37^.

### Plastic and evolved division of metabolic labour between parents and offspring

Regardless of their experimental evolutionary history, when an adult was not there to provide post-hatching care, larvae were able to produce many oral proteins for themselves. This plastic response to a lack of parental assistance could be an adaptation for the highly variable duration of care seen in natural populations, including (occasionally) no post-hatching care at all^11,16^. Furthermore, this plasticity is probably regulated by a simple cue, such as the presence or absence of a parent, because we found that its fine-tuning was somewhat limited. When mothers were allowed to tend to their offspring, the protein secretions in each brood’s oral fluids were not clearly attuned to the protein secretions produced by their mother. Future work pairing both transcriptomics and proteomics would allow for more clear testing of these possibilities.

Although socially induced plasticity in protein production accounted for more of the variation in larval oral fluid proteins, we found some evidence of evolutionary divergence in protein production between No Care and Full Care populations. These differences were most apparent when larvae from both the No Care and Full Care populations were raised without post-hatching care. Thus, rapid microevolutionary change involved a change in the extent to which protein production could be socially modulated rather than a change in the proteins secreted, which is consistent with ‘plasticity-first’ models of evolutionary change^38^. When parents were absent, the five proteins that significantly differed in the larval fluids between the evolutionary populations were all connected to immunity functions. Each could potentially link directly to larval fitness, by facilitating superior immune defences in larvae against the rich microbial environment on the carrion nest. We infer that No Care larvae adapted to the No Care environment by more readily expressing oral fluid proteins that could promote their survival without parents. Although we found evidence that larvae from the No Care populations were able to consume more mouse protein when developing without parents than were larvae from the Full Care populations, there was no associated difference in the digestive proteins in their oral fluids suggesting that differences in mouse consumption were instead mediated by differences in microbial control, or possibly through other means not tested here (e.g. behaviour, non-protein molecules).

In conclusion, by focusing on the proteins transferred orally within burying beetle families, we have been able to see in detail how parents share some of the metabolic burden of offspring development, both with each other and with their brood. We have shown that the production of oral fluid proteins can be plastically modulated, and that the extent of plasticity is a target for microevolution when social conditions change. The limits on the extent of this plasticity appear to account for the extent to which larvae are dependent on parents to survive.

## Methods

### Study taxon

Burying beetles *Nicrophorus vespilloides* vary between families in the extent of biparental care they supply^16^. Pairs convert a small dead vertebrate into an edible nest for their larvae. Once they have located a carcass they tear off any fur or feathers, spread it with anal and oral fluids, roll it into a ball and bury it underground^6^. Eggs are laid in the surrounding soil during this time. After hatching, larvae crawl to the carrion nest where they are tended by their parents. If parents die or desert following nest preparation then larvae can self-feed on the carrion and survive to become adults^39^.

### Experimental Evolution

The *N. vespilloides* populations used in this study were part of a long-term experimental evolution project that investigated how populations of burying beetles adapt to different regimes of parental care. These populations were founded in 2014 with wild-caught beetles from four woodland sites across Cambridgeshire, UK (Byron’s Pool, Gamlingay Woods, Waresley Woods and Overhall Grove). Full details can be found in^19^. The populations were ceased in 2021.

There were four experimental populations: ‘Full Care’ (x2 replicates) and ‘No Care’ (x2 replicates). Full Care populations were allowed to prepare the carcass nest and stay with offspring until dispersal, as described above. However, in the No Care populations at 53 h post-pairing, when the carrion nest was complete and eggs were laid, but before the larvae had hatched, both parents were removed to prevent post-hatching care. This procedure was repeated at every generation. The life cycle of beetles within each replicate block (Full Care 1/No Care 1 and Full Care 2/No Care 2) was staged to run 7 days apart.

### Burying beetle husbandry in the laboratory

Each pair of sexually mature unrelated male and female beetles was bred in a plastic breeding box (17 x 12 x 6 cm) with damp soil (Miracle-Gro Compost) and a 10-15 g mouse carcass, kept in constant darkness at 21°C. At natural dispersal time (8 days after pairing) larvae were counted, weighed and placed in plastic pupation boxes (10 x 10 x 2 cm) filled with damp peat. Juvenile adults were eclosed approximately 21 days later and housed in individual boxes (12 x 8 x 2 cm). Adults were fed twice a week (beef mince) until breeding at 15 days post-eclosion (when sexually mature). Both adult and pupating larvae were kept on a 16L: 8D hour light cycle at 21°C.

### Experimental breeding design

In preparation for this experiment, at generations 49 and 50 beetles from all populations (including No Care 1 and No Care 2) were put through two generations of a common garden Full Care environment treatment to control for any potential gene by environment effects^40^. We collected samples from beetles breeding in all four populations in generation 51.

### Fluid collection

Oral fluids were collected by holding each beetle or larva by the abdomen and rubbing a 5 ul glass microcapillary tube (Drummond) along its mouth until it regurgitated fluid (supplementary video 1). Each sample was pipetted immediately into 1.5 ml Eppendorf tubes containing 5 ul of 1 x Sigmafast Protease Inhibitor Cocktail (Sigma-Aldrich) on dry ice and subsequently stored at −80 °C until analysis.

#### Pre-hatching sampling

We collected oral fluids at 53 h after pairing (i.e. at the end of carcass preparation but before egg hatching) from both parents in *n* = 8 No Care and *n* = 8 Full Care breeding pairs. In one pair from each treatment one of the parents did not produce sufficient oral fluids for analysis, resulting in a sample size of *n* = 7 pairs for each treatment for the within-pair comparisons.

#### Post-hatching sampling

Pairs that were not sampled at pre-hatching were assigned to one of four treatments:

1. Full Care pairs allowed to provide full post-hatching care = FC_POP_ FC_ENV_ (Block 1, *n* = 14; Block 2, *n* = 11)
2. Full Care pairs which were removed at 53 h after pairing to prevent post-hatching care = FC_POP_ NC_ENV_ (Block 1, *n* = 14; Block 2, *n* = 17)
3. No Care pairs which were removed at 53 h after pairing to prevent post-hatching care = NC_POP_ NC_ENV_ (Block 1, *n* = 16; Block 2, *n* = 17)
4. No Care pairs allowed to provide full post-hatching care = NC_POP_ FC_ENV_ (Block 1, *n* = 16; Block 2, *n* = 18)

This full factorial design allowed us to determine whether any differences between the populations, in either the larvae or the parents, were plastically induced by the care environment they were currently experiencing or due to evolutionary divergence between populations.

At the start of the third instar of larval development (four days after pairing for FC_POP_ FC_ENV_ and NC_POP_ FC_ENV_, and five days after pairing for FC_POP_ NC_ENV_ and NC_POP_ NC_ENV_), oral fluids were collected from nine larvae in every brood and pooled. This larval stage of development was chosen for sampling because the larvae were large enough to reliably sample but were still likely to be receiving some fluids by trophallaxis. Broods with fewer than nine larvae at this stage were not sampled to ensure standardisation. Oral fluids were also collected from the mothers of broods where parents were allowed to provide post-hatching care (FC_POP_ FC_ENV_ and NC_POP_ FC_ENV_). Mothers were chosen for sampling because we found fathers did not produce sufficient oral fluids to collect. Of the pairs we bred, the following numbers had at least nine surviving third instar larvae. Broods in the FC_POP_ NC_ENV_ condition performed poorly, resulting in a lower sample size:

1. FC_POP_ FC_ENV_ = Block 1: 13 of 14 pairs, Block 2: 10 of 11 pairs
2. FC_POP_ NC_ENV_ = Block 1: 3 of 14 pairs, Block 2: 5 of 17 pairs
3. NC_POP_ NC_ENV_ = Block 1: 10 of 16 pairs, Block 2: 10 of 16 pairs
4. NC_POP_ FC_ENV_ = Block 1: 15 of 16 pairs, Block 2: 12 of 18 pairs

### Proteomics

Oral fluid samples (supplementary table 6) were processed in two batches (the post-hatching time point samples and two samples from the pre-hatching time point in January 2021, and the remaining pre-hatching time point samples in January 2022) as in^23^, and LC-MS/MS measurements were performed following the same protocols on a QExactive plus mass spectrometer (Thermo Scientific) coupled to an EasyLC 1000 nanoflow-HPLC. The haemolymph samples were run together with the January 2021 batch, and used to assess oral fluid sample purity.

The MS raw data files were uploaded into MaxQuant software^41^, version 1.6.2.10, for peak detection, generation of peak lists of mass error corrected peptides, and for database searches. To annotate the proteins, MaxQuant was set up to search the reference genome of *N. vespilloides* (NCBI GCF_001412225.1), along with common contaminants and the Uniprot *Mus musculus* database (April 2016). Carbamidomethylcysteine was set as fixed modification and protein amino-terminal acetylation and oxidation of methionine as variable modifications. Three missed cleavages were allowed, enzyme specificity was trypsin/P, and the MS/MS tolerance was 20 ppm. Peptide lists were further used by MaxQuant to identify and relatively quantify proteins using the following parameters: peptide and protein false discovery rates, based on a forward-reverse database, were set to 0.01, minimum peptide length was set to 7, minimum number of peptides for identification and quantitation of proteins was set to one which must be unique. The ‘match-between-run’ option (0.7 min) was used. For the protein abundance comparisons among sample types, all proteins labelled as contaminants, reverse or only identified by site were excluded, as were all mouse proteins, and only proteins with scores higher than 70 were used. After these filtering steps, the dataset contained 200 *N. vespilloides* proteins.

### Statistical analyses on beetle protein abundances

Quantitative analysis was performed in Perseus version 1.6.15.0^41^ using iBAQ values^42^. The full dataset was divided into subsets and within each subset only proteins present in over 70% of the samples were analysed, to filter out proteins present predominantly in the haemolymph samples and the other subsets. The subsets comprised: a) parents at the pre-hatching time point (53 h): for analysing the difference between males and females, and the difference between the evolutionary populations (123 proteins); b) females at the post-hatching time point (third larval instar of development): for analysing the difference between evolutionary populations (142 proteins); and c) larvae at the post-hatching time point: for analysing the difference between evolutionary populations and the effect of parents’ presence or absence (122 proteins). As the parents’ presence (current care regime) had such a strong effect at the post-hatching time point, and there was likely transfer from the parents when they are present, the larval dataset was further divided based on the current care regime, and the effect of the experimental population was analysed separately for both. For the pre-hatching time point samples, the two samples whose LC-MS/MS measurements were run in a different batch (together with post-hatching time point samples), separated from the rest due to a technical difference in overall abundances, and were thus excluded. No analyses were run across the time points due to this batch effect.

All above comparisons were run as two-sample t-tests, with log2 transformed and median-centred data, with missing data imputed by random sampling from normal distribution with 2SD downward shift and 0.3 width for each sample. Permutation-based FDR of 0.05, and S0 parameter was set to two for all analyses.

Euclidean distance between samples was computed with MATLAB R2020b using the pdist function for a) male-female pairs at the pre-hatching time point, and b) mother-offspring pairs at the post-hatching time point. The greater the Euclidean distance, the more different we inferred the protein content of the samples to be. Thus, the Euclidean distance should be small when social partners are producing very similar proteins in their oral fluids and greater when the proteins produced by each social partner are complementary. Linear models were fitted in R, gamma-distributed for a) and Gaussian for b), to analyse whether distance differs between false pairs and true pairs, and between the evolutionary populations. Also, the experimental population of the mothers (Full Care/No Care 1 or Full Care/No Care 2) was included in the model to account for the non-independence of the samples. To infer whether there was relaxed selection in the protein abundances in the No Care females by analysing variation in protein abundance, proteome variability was compared between the Full Care and No Care evolutionary populations at the post-hatching time point, in R with a linear model of the square-rooted coefficient of variation of the iBAQ abundance of each protein.

### Beetle and mouse protein proportion analyses

To analyse the relative abundance of endogenously produced beetle proteins vs. exogenous mouse proteins in the oral fluid samples, the sums of intensities of all proteins were calculated for each protein class and each sample. The intensity was used here instead of iBAQ, because it better represents the total mass of protein. The proportion of beetle produced proteins was compared against a hypothetical 50/50 distribution, separately for adults and offspring samples, with a one-sample t-test. The proportion of beetle-produced proteins was compared among the different sample types by fitting a linear model on the data in R version 4.3.1^43^. No significant effect of block on protein proportions was detected - except for one treatment: FC_POP_ NC_ENV_ larvae. However, in this condition the larvae performed overall more poorly than in the others, yielding *n* = 3 in block 1 and *n* = 2 in block 2. Since robust conclusions cannot be derived from such a small sample size, samples from the two blocks were analysed together.

### Protein annotation and gene set enrichment analysis

For annotations, all proteins were protein BLASTed against the *Drosophila melanogaster* genome. The protein functions were surveyed from Flybase.org, and all proteins were classified into one or more of following categories: Growth (*n* = 36), Digestion (*n* = 73), Immunity (*n* = 25), Communication (*n* = 13), Other/Unknown (*n* = 49).

GO analysis and protein-protein interaction and pathway analyses were performed with STRING version 11.5^44^. We used *D. melanogaster* orthologues of the 65 most abundant proteins, separately for all the samples of the pre-hatching time point and post-hatching time point (pre-hatching time point, *n* = 56 of the proteins that had corresponding *Drosophila* gene names, and post-hatching time point *n* = 57). Similar analyses were run for the proteins (*n* = 44) that significantly differed in any of the analyses, since they were most likely to have functional roles that could vary with the experimental treatments.

## Data availability

Data have been uploaded to ProteomeXchange and an accession number will be provided when acquired.

## Acknowledgements

E.K.B. was supported by a Biotechnology and Biological Sciences Research Council PhD studentship (BB/M011194/1). S.M.H was supported by a grant from the Finnish Cultural Foundation. R.M.K. was supported by a Consolidator’s Grant from the European Research Council (310785 Baldwinian_Beetles), a Wolfson Merit Award from the Royal Society, The Leverhulme Trust (RPG-2018-232), and The Isaac Newton Trust (18.23(q)). A.C.L was supported by a Swiss National Science Foundation PRIMA grant PR00P3_179776.

We thank Benjamin Jarrett, Darren Rebar, Matthew Schrader and Rahia Mashoodh for establishing and helping to maintain the experimental evolution populations, and Chris Swannack and Sue Aspinall for beetle maintenance during the initial generations of experimental evolution. We thank Michael Stumpe from the Metabolomics and Proteomics Platform at the University of Fribourg for his work and support.

