## Supplementary figures for "Plasticity and evolution of metabolic division of labour within families"

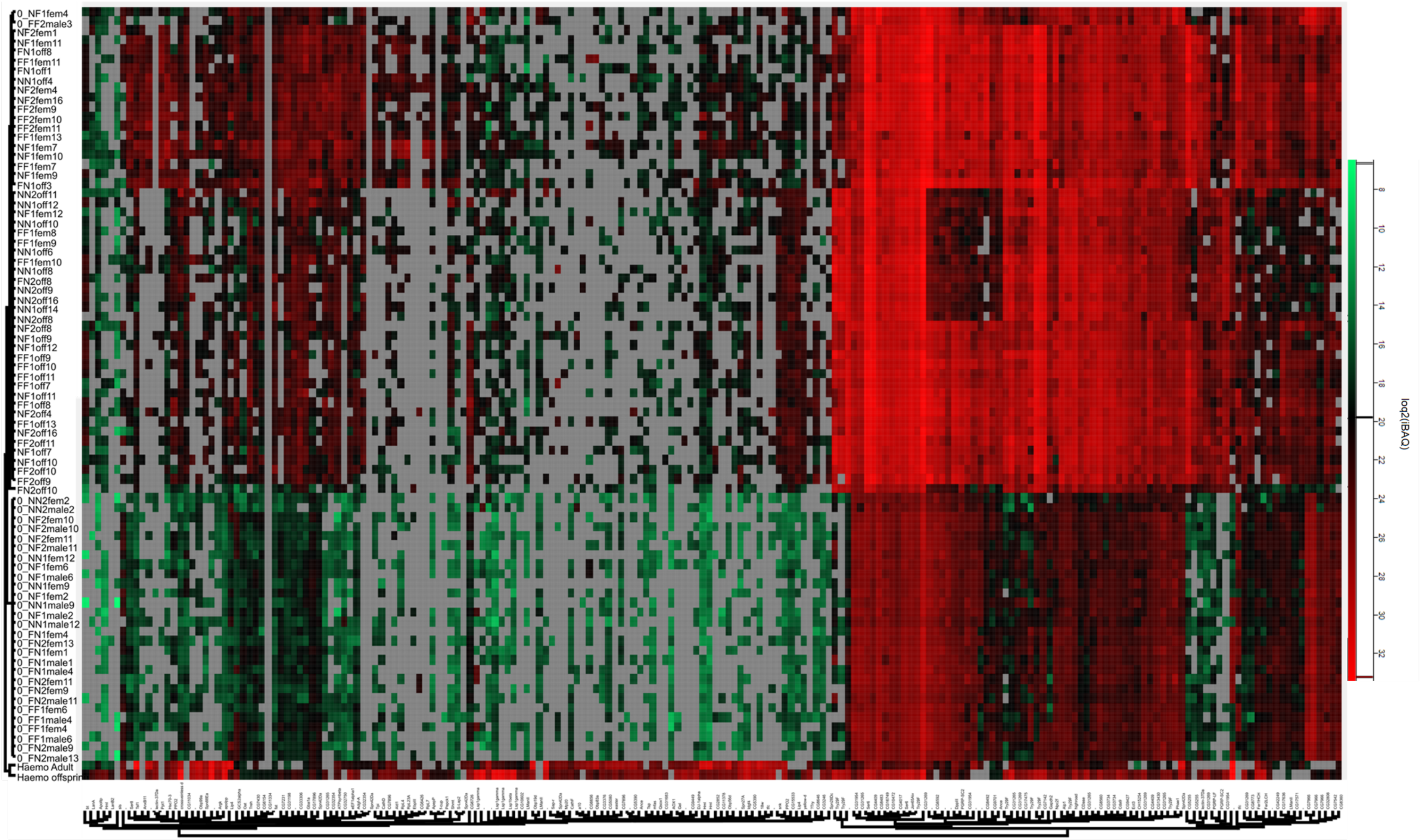


Figure S1: Heatmap of proteins present in samples, based on log2 (iBAQ) values. Grey indicates a protein is missing from a sample (or is at too low a quantity to be detected), green that it is in low abundance and red that it is in high abundance.

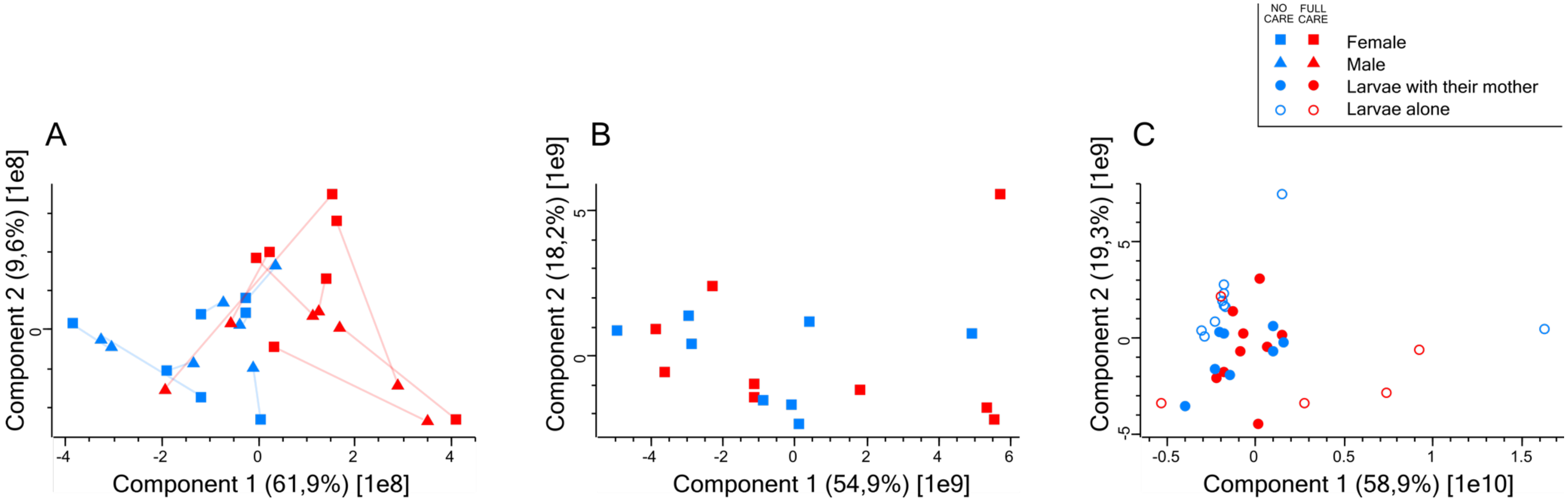


Figure S2: Principal components analyses for A) male and female oral fluids at the pre-hatching time point with pairs connected with lines, B) oral fluids of females providing care ($NC_{POP}FC_{ENV}$ or $FC_{POP}FC_{ENV}$) at the post-hatching time point in the experimental generation, and C) larval oral fluids with and without post-hatching care in the experimental generation (all treatments). Red = Full Care evolutionary populations, blue = No Care evolutionary populations. Squares = females, triangles = males, full circle = offspring with post-hatching care, open circle = offspring with no post-hatching care.
